# Heart-Brain Interactions Shape Somatosensory Perception and Evoked Potentials

**DOI:** 10.1101/750315

**Authors:** Esra Al, Fivos Iliopoulos, Norman Forschack, Till Nierhaus, Martin Grund, Paweł Motyka, Michael Gaebler, Vadim V. Nikulin, Arno Villringer

## Abstract

Human perception either refers to the external world, exteroception, or internal body parts such as the heart, interoception. How these two types of perception interact is poorly understood. Using electroencephalography, we identify two heartbeat-related modulations of conscious somatosensory perception: (i) When stimulus timing coincided with systole of the cardiac cycle, participants were less likely to detect and localize somatosensory stimuli, and late components (P300) of the somatosensory-evoked potential (SEP) were attenuated. (ii) The amplitude of the heartbeat-evoked potential (HEP) negatively correlated with detection bias (criterion) and localization accuracy. Furthermore, higher HEP amplitudes were followed by decreases in both early and late SEP amplitudes. Both heartbeat-related effects were independent of the alpha oscillations’ influence on somatosensory processing. We conclude that internal signals are integrated into our conscious perception of the world, and connect our results to predictive processing (heartbeat-coupled stimulus timing) and attentional shifts between exteroception and interoception (HEP amplitude).

## INTRODUCTION

The neural response to an external stimulus and its access to consciousness depend on stimulus features as well as the state of the brain (Arieli, Sterkin, Grinvald, & Aertsen, 1996; Blankenburg et al., 2003; Gelbard-Sagiv, Mudrik, Hill, Koch, & Fried, 2018; van Dijk, Schoffelen, Oostenveld, & Jensen, 2008; Weisz et al., 2014). Interestingly, functional states of other bodily organs, such as the heart, can also influence the perception of external stimuli. For example, timing along the cardiac cycle (e.g., systole versus diastole) can impact the perception of visual, auditory, and somatosensory stimuli (Motyka et al., 2019; Salomon et al., 2016; Sandman, McCanne, Kaiser, & Diamond, 1977; Saxon, 1970). Likewise, neural responses to visual and auditory stimuli are modulated across the cardiac cycle (Sandman, 1984; Walker & Sandman, 1982). Most often they have been reported to be higher during diastole than systole (Sandman, 1984; Walker & Sandman, 1982). A recent study (Park, Correia, Ducorps, & Tallon-Baudry, 2014) has also associated fluctuations of the heartbeat-evoked potential (HEP; Kern, Aertsen, Schulze-Bonhage, & Ball, 2013; Montoya, Schandry, & Müller, 1993; Pollatos & Schandry, 2004) with conscious detection of a visual stimulus.

While thus increasing evidence indicates that events related to cardiac function may modulate conscious perception, fundamental questions remained unanswered. Is it perceptual discrimination ability, i.e., *sensitivity* in signal detection theory (SDT; Green & Swets, 1966), that is influenced by cardiac activity? Or, might a bias to report the presence or absence of a stimulus underlie the effect, i.e., *criterion* in SDT? Are criterion-free decisions also affected by the heart? How are these perceptual effects reflected in evoked neural activity? More specifically, do these effects influence early, preconscious, somatosensory-evoked potentials (SEP), or, only the late components? Ultimately, how cardiac-related modulation of perceptual awareness relates to primary determinants of sensory perception and evoked brain activity, such as *prediction*, *attention*, and *background neural activity*, is unknown.

The current study targets mechanisms linking heart, brain, and perception using a somatosensory detection and localization task with electroencephalography (EEG) recordings. In an SDT-based design, we identify differential effects of two heartbeat-related phenomena, i.e., (i) stimulus timing during the cardiac cycle and (ii) the amplitude of the HEP, on somatosensory perception and evoked potentials. We argue that these findings are in line with a predictive coding account for cardiac phase-related sensory fluctuations and spontaneous shifts between interoception and exteroception as indexed by the HEP amplitude.

## RESULTS

### Behavior

Thirty seven participants were presented weak somatosensory (electrical) stimuli to either the left index or middle finger in a combined *yes/no detection* and *location discrimination* task (**Fig. 1**). Both EEG and electrocardiography (ECG) were recorded. On average, participants detected 51.0 ± 10.5% (mean ± s.d.) of the somatosensory stimuli with a false alarm rate of 8.4 ± 7.7%. This corresponds to a mean detection sensitivity, d’, of 1.57 ± 0.57 and a decision criterion, c, of 0.76 ± 0.32. Participants correctly localized 73.3 ± 6.6% of stimuli (finger-wise), corresponding to a mean localization sensitivity of 0.90 ± 0.32. Participants correctly localized 88.9 ± 7.9% of hits and 57.0 ± 6.9% of misses.

**Figure 1.**
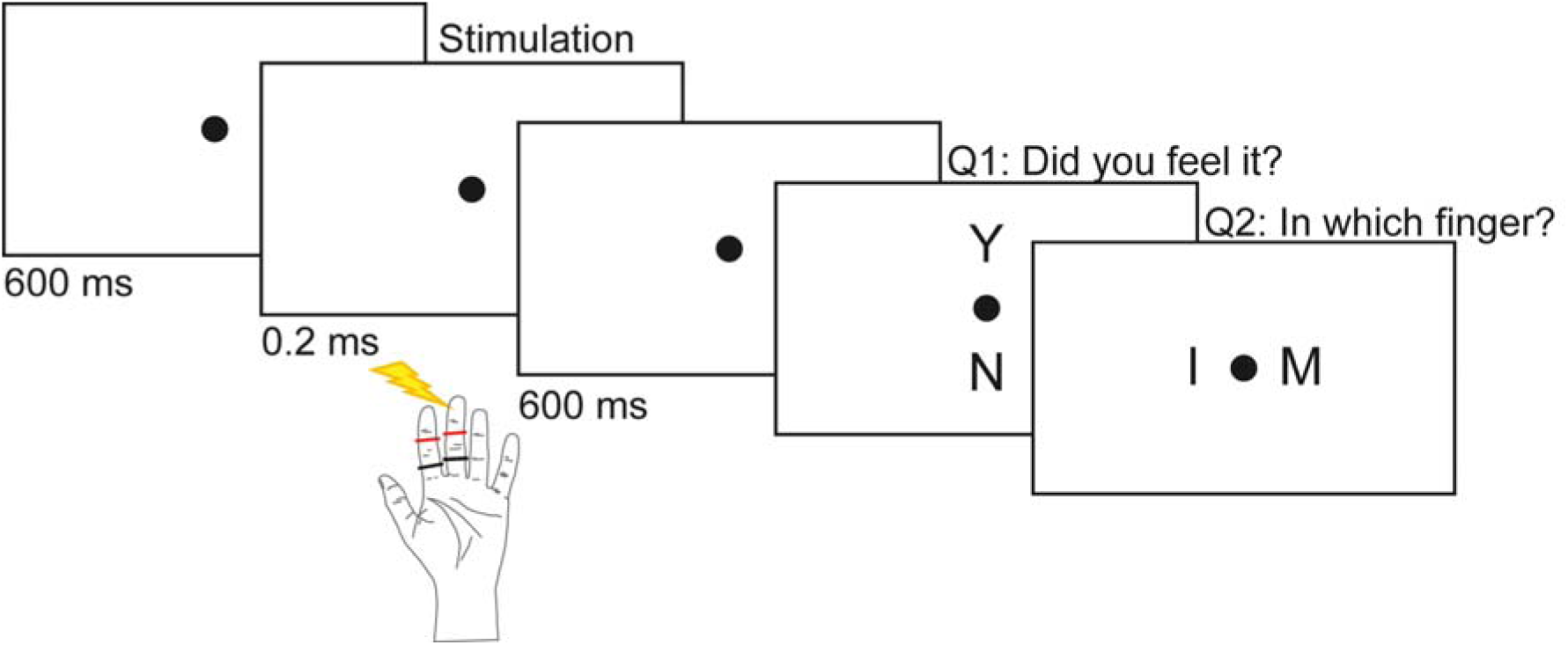
Experimental paradigm. 37 subjects received a weak electrical pulse to the left index or the middle finger in 800 out of 960 trials over 8 experimental blocks. Subjects were told that every trial contained a stimulus, however, in 160 pseudo-randomized trials no stimulus was actually presented. In every trial, participants were asked to first perform a yes/no detection task and then a location discrimination task.

### Detection varies across the cardiac cycle

We hypothesized that hits were more likely to occur in a later phase of the cardiac cycle, whereas misses would occur in an earlier phase (Motyka et al., 2019). A Rayleigh test showed that hits were not uniformly distributed, 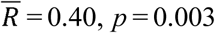 (**Fig. 2a**), with a mean angle of 308.70° corresponding to the later cardiac cycle phase (i.e., diastole). Similarly, the distribution of misses was not uniform, 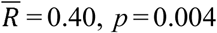 (**Fig. 2a**), with a mean angle of 93.84°, located in the early phase of the cardiac cycle (i.e., systole). We observed a trend in the distribution of correct localizations towards the later phases of the cardiac cycle (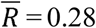, *p* = 0.067). The distribution of wrong localizations was not significantly different from a uniform distribution, 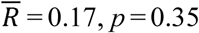 (**Fig. 2a**).

**Figure 2.**
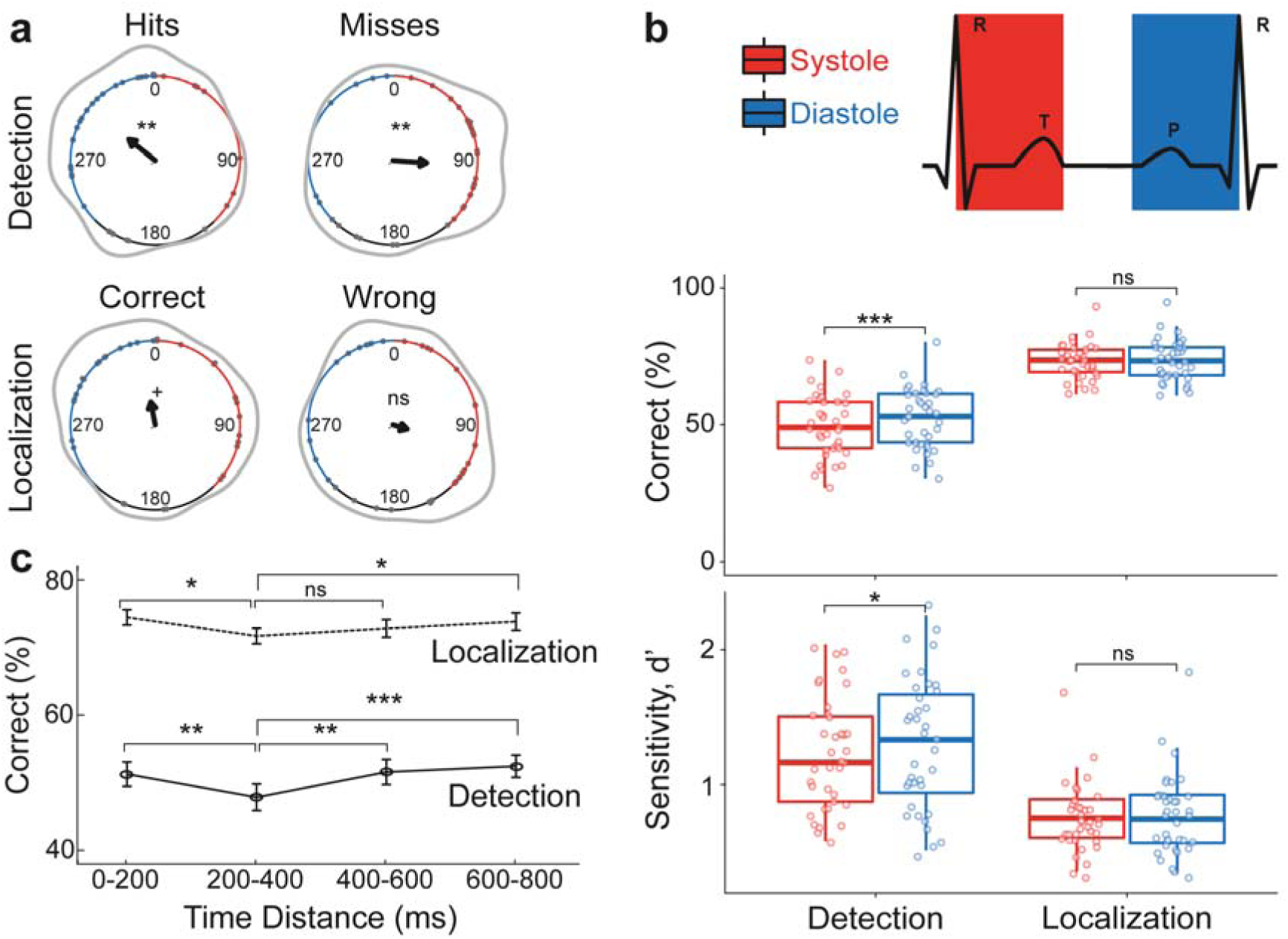
Conscious detection of somatosensory stimuli varies across the cardiac cycle. (**a**) Distribution of hits (*top-left*), misses (*top-right*), correct localizations (*bottom-left*) and wrong localizations (*bottom-right*) across the cardiac cycle (the interval between two R-peaks at 0/360°). Gray points show subjects’ mean degrees. The black arrows point towards the overall mean degree and its length indicates the coherence of individual means. The gray lines depict the circular density of individual means. The overall mean systole and diastole lengths are shown with red and blue, respectively. Hits and misses were non-uniformly distributed across the cardiac cycle (Rayleigh tests, 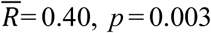 and 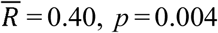, respectively). While correct localizations showed a trend towards a non-uniform distribution (*p* = 0.067), wrong localizations did not show a significant deviation from uniform distribution (*p* = 0.35). (**b**) *Top*: Correct detection and localization percentages during systole and diastole. Participants had more correct detections in diastole (*t*_36_ = −3.95, *p* = 3·10^−4^). No statistically significant difference between systole and diastole was found for correct localization (*p* = 0.54). *Bottom*: Detection and localization sensitivity (d’) between systole and diastole. Detection sensitivity was significantly higher in diastole than systole (*t*_36_ = −2.38, *p* = 0.008), localization sensitivity did not differ significantly between the two cardiac phases (*p* = 0.38). **c**) Correct detection and localization of somatosensory stimuli relative to their distance from the previous R-peak. Both detection and localization performances were lowest 200 – 400 ms after the R-peak. (*post-hoc* paired *t-*test between 0 – 200 and 200 – 400 ms for detection: *t*_36_ = 3.76, *p* = 6·10^−4^ and localization: *t*_36_ = 2.88, *p* = 0.007). Error bars represent SEMs. +*p* < 0.08, \**p* < 0.05, \*\**p* < 0.005, \*\*\**p* < 0.0005, ns, not significant.

### Detection rate and sensitivity are higher during diastole compared to systole

We also examined detection and localization performance by segmenting the cardiac cycle into two cardiac phases: systole and diastole. As suggested by our first analysis, the detection rate for the weak stimuli was significantly higher during diastole (M = 52.41 %) than systole (M = 49.53 %), *t*_36_ = −3.95, *p* = 3·10^−4^ (**Fig. 2b**). Increased detection rate during diastole was observed for 27 out of 37 participants. However, the false alarm rate did not differ significantly between systole (M = 8.50 %) and diastole (M = 8.19 %), *t*_36_ = 0.54, *p* = 0.59. We furthermore tested whether the effect of cardiac phase on detection correlated with the heart rate or the heart rate variability (i.e., the standard deviation of RR intervals) of individuals. While there was no significant correlation between subject’s heart rate and their detection rate variation between systole and diastole (Pearson’s correlation, *r* = 0.01, *p* = 0.95), subjects’ heart rate variability negatively correlated with their detection rate difference (*r* = −0.36, *p* = 0.03, **Supplementary Fig. 1**).

Signal Detection Theory (SDT) was applied to test whether the increased detection rates in diastole were due to increased perceptual sensitivity (d’) or due to adopting a more liberal response strategy (criterion). Detection sensitivity was significantly higher in diastole (M = 1.59) than systole (M = 1.48), *t*_36_ = −2.38, *p* = 0.008 (**Fig. 2b**). For the criterion, no significant difference between systole (M = 0.75) and diastole (M = 0.73) was found *t*_36_ = 0.71, *p* = 0.48. Localization performance was also tested across the cardiac cycle. Correct localization rate did not differ significantly between systole (M = 73.27 %) and diastole (M = 73.68 %), *t*_36_ = −0.62, *p* = 0.54. Likewise, localization sensitivity was not significantly different between systole (M = 0.90) and diastole (M = 0.93), *t*_36_ = −0.89, *p* = 0.38 (**Fig. 2b**).

In an exploratory analysis, we tested whether detection and localization rates differed as a function of the distance of stimulus onset from the previous R-peak, in four time windows: 0 – 200, 200 – 400, 400 – 600, and 600 – 800 ms. Detection and localization rates were significantly different between these time windows (within-subject ANOVA, *F*_3,108_ = 7.25, *p* = 2·10^−4^ and *F*_3,108_ = 3.97, *p* = 0.01). Detection and localization was lowest 200 – 400 ms after the R-peak (*post-hoc* paired *t-*test between 0 – 200 and 200 – 400 ms windows for detection: *t*_36_ = 3.76, *p* = 6·10^−4^ and localization: *t*_36_ = 2.88, *p* = 0.007; between 200 – 400 and 400 – 600 ms for detection: *t*_36_ = − 3.61, *p* = 9·10^−4^ and localization: *t*_36_ = −1.36, *p* = 0.18; **Fig. 2c**). Significant differences were found for the sensitivity (main effect of time, *F*_3,108_= 6.26, *p* = 6·10^−4^; *post-hoc* paired *t-*test between 0 – 200 and 200 – 400 ms, *t*_36_ = 2.83, *p* = 0.008 and between 200 – 400 and 400 – 600 ms, *t*_36_ = −3.48, *p* = 0.001) but not for the criterion (*F*_3,108_ = 0.10, *p* = 0.96; **Supplementary Fig. 2**).

### Somatosensory-evoked potentials during diastole compared to systole

Conscious somatosensory perception is known to correlate with greater amplitude of certain somatosensory-evoked potential (SEP) components such as N140 and P300 (Auksztulewicz & Blankenburg, 2013). In line with the changes in somatosensory perception, we expected to find differences in SEPs during diastole compared to systole. We systematically compared SEPs during systole and diastole in the time window of 0 (stimulation onset) – 600 ms with a cluster-based permutation *t*-test. SEPs over the contralateral somatosensory cortex (indexed by C4 electrode) showed greater positivity when stimulation was performed during diastole than systole in two temporal clusters: 268 – 340 ms and 392 – 468 ms, (Monte Carlo *p* = 0.004 and *p* = 0.003, respectively, corrected for multiple comparisons in time; **Fig. 3a**). SEPs for hits during diastole and systole did not differ significantly (smallest Monte-Carlo *p* = 0.27). SEPs for misses, however, differed between systole and diastole over the contralateral somatosensory area. Higher positivity was observed in diastole compared to systole in time windows of 288 – 324 ms and 400 – 448 ms, respectively (Monte-Carlo *p* = 0.02 and Monte-Carlo *p* = 0.01, respectively; **Fig. 3c**).

**Figure 3.**
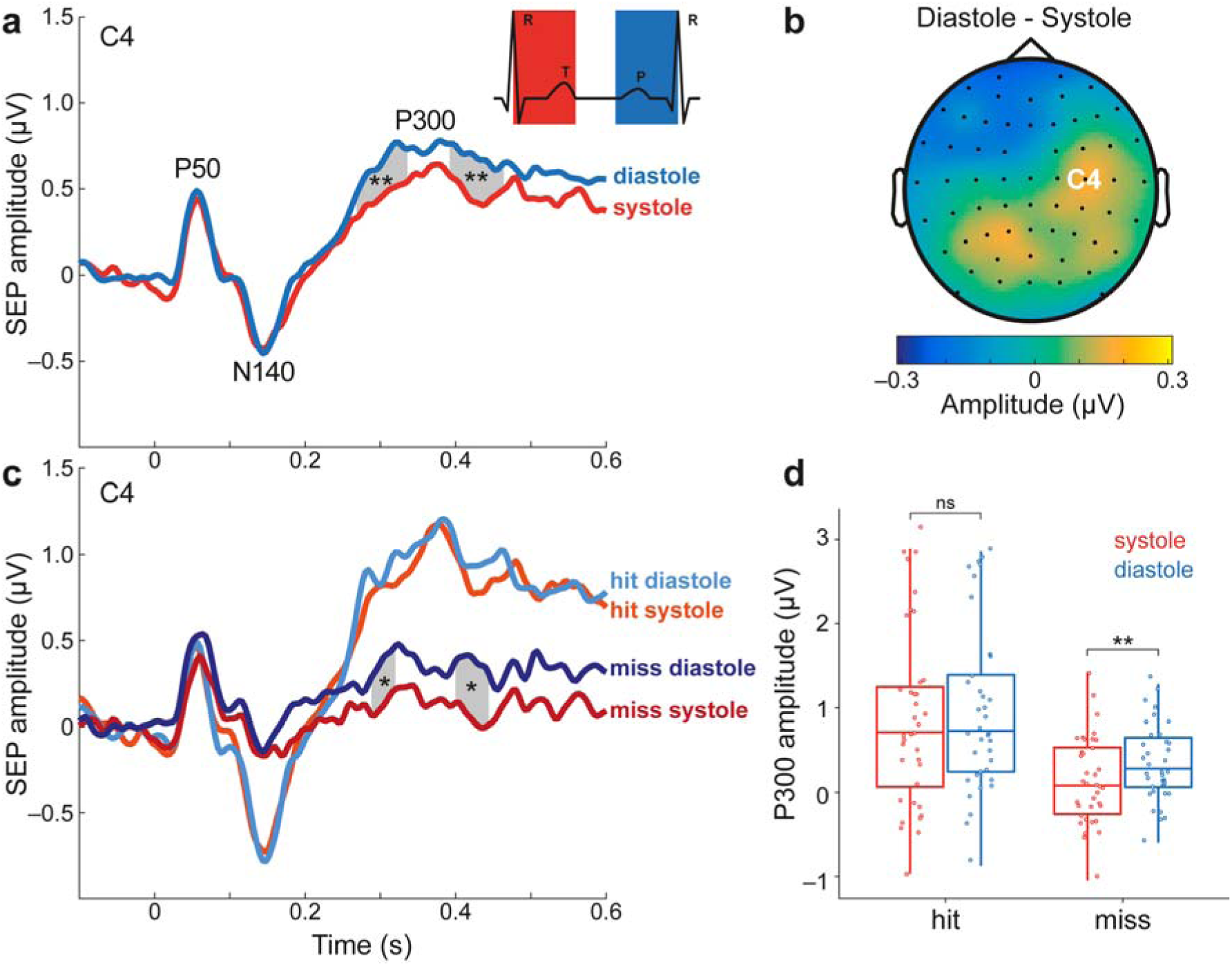
Somatosensory-evoked potentials (SEPs) for stimulations during systole versus diastole (**a**) The difference in P300 component of SEPs (electrode C4) between systole and diastole. SEPs were more positive for stimuli during diastole than systole between 268 – 340 ms and 392 – 468 ms after stimulus onset over contralateral somatosensory cortex (Monte Carlo *p* = 0.004 and *p* = 0.003, respectively, corrected for multiple comparisons in time). (**b**) The topography contrast between diastole and systole between 268 and 468 ms. The position of electrode C4 is shown on the head model (**c**) SEPs for hits (lighter colors) and misses (darker colors) during systole (red) and diastole (blue). SEPs showed higher positivity for misses during diastole than during systole in two time windows: 288– 324ms and 400–448 ms (*p* = 0.02 and *p* = 0.01, respectively). (**d**) The mean SEP amplitude between 268–468 ms for detection and cardiac phases. \**p* < 0.05, \*\**p* < 0.005, ns, not significant.

We used a within-subject ANOVA with the factors: detection (hit vs. miss) and cardiac phase (systole vs. diastole) to examine their effect on the P300 component of the SEPs. The P300 latency was determined in the 268 – 468 ms interval by merging the two time clusters observed for SEP differences between systole and diastole. We found significant main effects of detection (F_1,36_ = 33.29, *p* = 1·10^−6^) and cardiac phase (F_1,36_ = 8.26, *p* = 0.007). We did not observe a significant interaction effect (F_1,36_ = 2.55, *p* = 0.12).

### Heartbeat-evoked potentials predict somatosensory detection

Heartbeat-evoked potentials (HEPs) are cortical electrophysiological responses time-locked to the R-peak of the ECG and are thought to represent neural processing of cardiac activity (Kern et al., 2013; Schandry, Sparrer, & Weitkunat, 1986; Schandry & Weitkunat, 1990). We tested whether HEPs immediately preceding stimulus onset, predicted somatosensory detection. To ensure that the time window for the HEP, *250 – 400ms after the R-peak*, (Kern et al., 2013; Schandry et al., 1986; Schandry & Weitkunat, 1990) was free of neural responses to the stimulation, we only included trials where the stimulus occurred at least 400 ms after the preceding R-peak (i.e., during diastole). We averaged the EEG data locked to the R-peak separately for hits and misses, and submitted the *250 – 40*0 ms *post R-peak time window* to a cluster-based permutation *t-*test. Prestimulus HEPs significantly differed between hits and misses over the contralateral somatosensory and central electrodes between 296 and 400 ms (Monte-Carlo *p* = 0.004 corrected for multiple comparisons in space and time; **Fig. 4a, b**) with a significantly higher positivity for misses. No significant changes were found in neither heart rate nor heart rate variability between hits and misses included in the HEP analyses (*t*_36_ = 1.51, *p* = 0.14 and *t*_36_ = −0.61, *p* = 0.55, respectively). Therefore, the observed differences in HEPs cannot be attributed to changes in heart rate or heart rate variability between hits and misses(Park et al., 2014).

**Figure 4.**
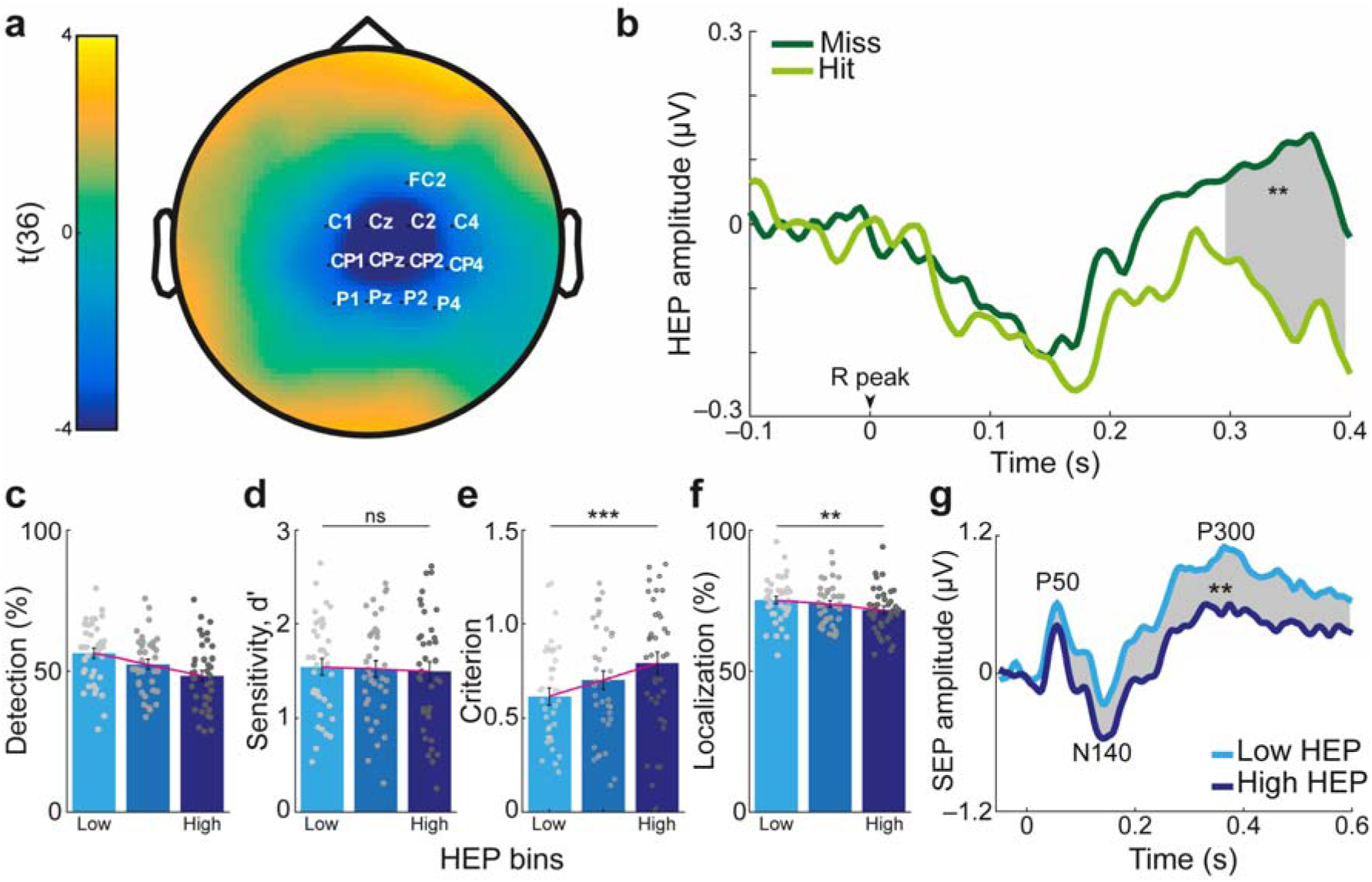
Heartbeat-evoked potentials (HEPs) before stimulus onset predicted somatosensory detection. (**a**) Topographical map of *t*-values for HEP differences preceding hits and misses: Grand average across 37 participants in the 296 – 400 ms time window, where a significant difference (misses>hits) was observed on the highlighted electrodes (Monte-Carlo *p* = 0.004 corrected for multiple comparisons in time and space). (**b**) Prestimulus HEPs averaged across the cluster. (**c-f**) Single-trials were sorted according to the mean HEP amplitude (across the cluster in the 296 – 400 ms time window) and split into three equal bins for each subject. (**c**) As the HEP amplitude increased, the detection rate decreased. (**d**) This decrease was not associated with a significant change in detection sensitivity (*p* = 0.84), (**e**) but correlated with an increase in criterion, i.e. reporting stimulus presence less often regardless of actual stimulus presence (p < 0.0005). (**f**) Similar to the decrease in detection rate, correct localization rate decreased with increasing HEP amplitude (*p* = 0.003). The gray points on the bar plots represent individual subjects. (**g**) Somatosensory evoked potential (SEP) amplitudes for trials in the low and high HEP bins. A significant difference in SEP amplitudes for the low and high HEP bin was observed between 32 – 600ms post-stimulation at contralateral somatosensory cortex (C4 electrode; Monte-Carlo *p* = 0.004 corrected for multiple comparisons in time). Error bars represent SEMs. \*\**p* < 0.005, \*\*\**p* < 0.0005, ns, not significant.

Subsequently, we calculated the prestimulus HEPs averaged across the cluster electrodes in the 296 – 400 ms time window separately for different detection responses (e.g., hits and misses). Similarly, we computed HEPs for cardiac cycles outside the stimulation window (see **Fig. 1**). Non-stimulation-related HEPs showed significantly more positivity than those preceding hits (paired *t-*test, *t*_36_ = 4.83, *p* = 3·10^−5^) and a trend towards more positivity compared to those preceding misses (paired *t-*test, *t*_36_ = 1.90, *p* = 0.07). HEP amplitudes preceding correct rejections showed significantly less positivity than HEPs preceding hits (paired *t-*test, *t*_36_ = 4.22, *p* = 2·10^−4^), and were not significantly different from HEPs preceding misses (paired *t-*test, *t*_36_ = 1.63, *p* = 0.11).

Next, we tested whether the HEP amplitude difference between hits and misses reflected a change in sensitivity or criterion according to SDT (**Fig. 4d-e**). We sorted single trials according to mean HEP amplitude (across the cluster electrodes in the 296 – 400ms time window) and split them into three equal bins (the number of HEP bins was chosen for comparability with a previous study(Park et al., 2014)) for each participant. We found that detection rates decreased as the HEP amplitude increased. Since we already showed this effect in the cluster statistics, we did not apply any statistical test here to avoid “double dipping”(Kriegeskorte, Simmons, Bellgowan, & Baker, 2009). The decrease in detection rate with increasing HEP amplitude was associated with an increase in criterion. More specifically, participants were more conservative in their decision and reported detecting the stimulus less often, regardless of their actual presence, when HEP amplitude was higher (within-subject ANOVA, *F*_2,36_ = 10.30, *p* = 1·10^−4^), Simultaneously, their sensitivity did not change significantly (*F*_2,72_ = 0.17, *p* = 0.84). We then tested whether prestimulus HEP amplitude could also affect somatosensory localization. Increasing HEP levels were associated with decreases in localization rate (*F*_1.72,62.01_ = 10.27, *p* = 0.03, **Fig. 4f**). Correct localization of hits and misses did not significantly differ between HEP bins (*F*_2,72_ = 1.26, *p* = 0.29 and *F*_2,72_ = 0.28, *p* = 0.76; **Supplementary Fig. 3**) indicating that the change in localization rate, associated with HEP amplitude, was connected with the change in detection rate.

We also tested whether prestimulus HEP amplitudes were associated with changes in SEP amplitudes. We applied a cluster-based permutation *t*-test in the time window of 0 – 600 ms (0 = stimulation onset) to compare SEPs following low and high HEP amplitudes. Between 32 ms and 600 ms SEPs over the contralateral somatosensory cortex had higher positivity when stimulation was preceded by low HEP compared to high HEP amplitudes (Monte-Carlo *p* = 0.004 corrected for multiple comparisons in time; **Fig. 4g**).

### Prestimulus sensorimotor alpha rhythm predicts somatosensory detection and localization

Given that alpha rhythm is known to influence sensory processing (Forschack, Nierhaus, Müller, & Villringer, 2017; Haegens, Nácher, Luna, Romo, & Jensen, 2011; Iemi, Chaumon, Crouzet, & Busch, 2017; Schubert, Haufe, Blankenburg, Villringer, & Curio, 2009; van Dijk et al., 2008), we assessed its effect on perception in our study as well as its possible interaction with heartbeat-related effects. Therefore, we sorted and divided trials into five equal bins (the number of alpha bins were chosen to be consistent with previous studies; Forschack et al., 2017; Iemi et al., 2017)), according to the mean sensorimotor alpha amplitude between 300 and 0 ms before stimulus onset. We then calculated the percentage of correct detection and localization responses for every bin. Correct detection and localization responses decreased with increasing levels of alpha amplitude (within-subject ANOVA, *F*_2.77,99.74_ = 8.88, *p* = 3·10^−7^ and *F*_3.30,118.81_ = 6.11, *p* = 4·10^−5^; **Fig. 5b**). With increasing prestimulus alpha amplitude, participants had a more conservative criterion (*F*_4,144_ = 3.77, *p* = 0.006; **Fig. 5c**). Sensitivity did not change significantly, but showed a trend toward a decrease (*F*_4,14_ = 2.20, *p* = 0.07; **Fig. 5c**).

**Figure 5.**
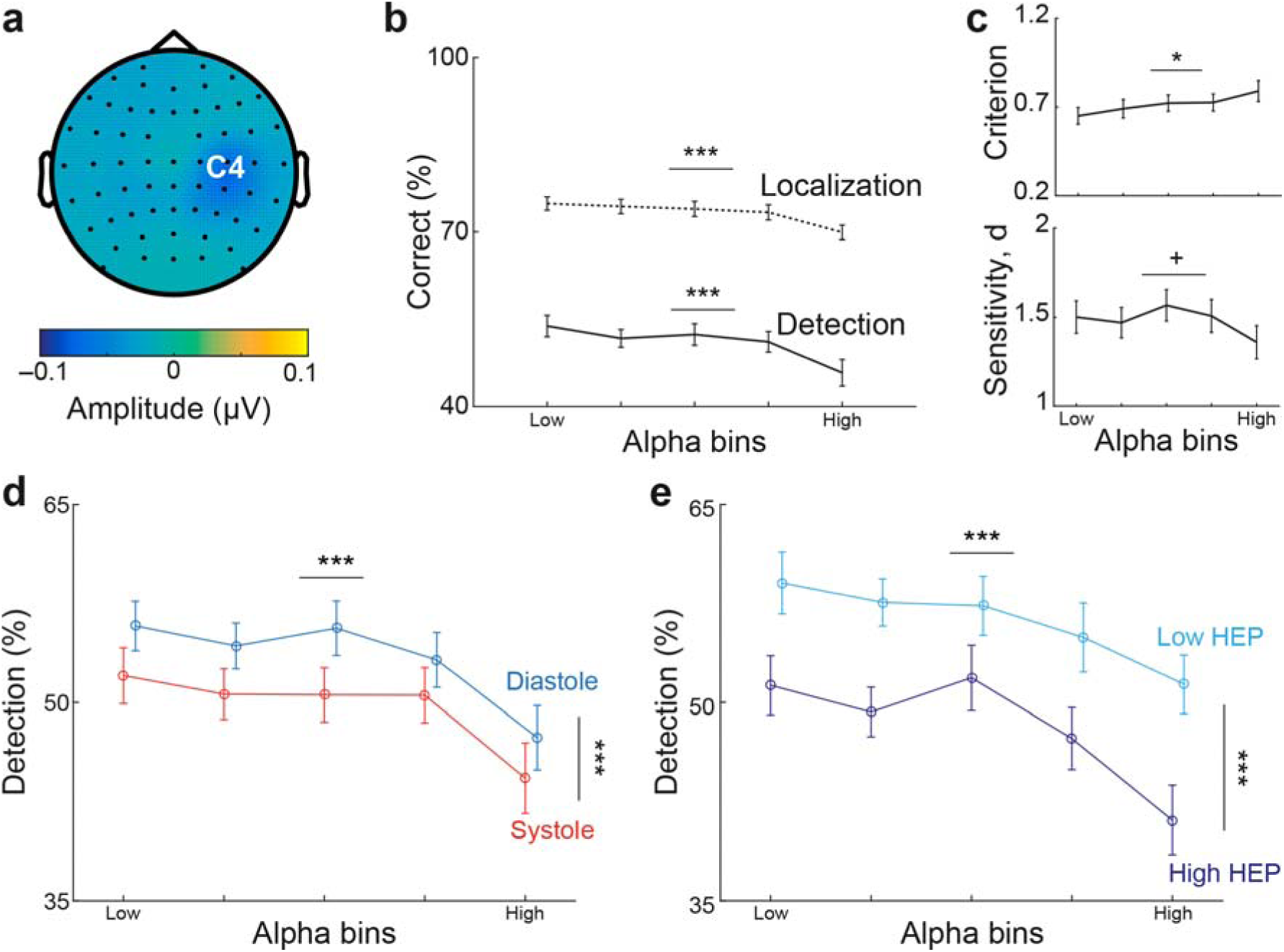
Prestimulus sensorimotor alpha amplitude affects somatosensory perception but does not mediate heartbeat-related perceptual effects. (**a**) Topography of prestimulus alpha (8 - 13 Hz) difference between hits and misses in the time window of 300 to 0 ms before stimulus onset. (**b**) Trials were sorted into five equal bins of increasing mean sensorimotor alpha amplitudes in the prestimulus time window of 300 to 0 ms over contralateral somatosensory cortex (C4 electrode). Correct detection and localization rates are given for each alpha bin. Both detection and localization decreased as alpha amplitude levels increased (*p* = 3·10^−7^ and *p* = 4·10^−5^). (**c**) The decrease in detection rates with increasing alpha amplitude levels was associated with a significant increase in criterion, i.e., a higher bias to miss the target (*p* = 0.006; top) and a trend towards lower sensitivity (*p* = 0.07; bottom). (**d**) For each alpha bin, detection rates are given separately for systole and diastole. Cardiac phase and alpha levels affected detection rate in an additive fashion (within-subject ANOVA test, *F*_1,36_ = 15.82, *p* = 3·10^−4^ and *F*_2.93,105.30_ = 12.05, *p* = 1·10^−6^). (**e**) For each alpha bin, detection rates are given separately for the trials with highest and lowest HEP, respectively. Prestimulus HEP amplitudes across the time window 296 – 400 ms after the R-peak were categorized in three equal bins for each participant, and detection rates were determined separately for the lowest and highest HEP conditions within each alpha bin. Both prestimulus factors, i.e., HEP amplitudes and alpha amplitudes influenced detection rates independently (within-subject ANOVA *F*_1,36_ = 38.71, *p* = 4·10^−7^ and *F*_4,144_ = 10.37, *p* = 2·10^−7^). Error bars represent SEMs. +*p* < 0.08, \**p* < 0.05, \*\*\**p* < 0.0005.

### Sensorimotor alpha does not mediate cardiac phase effect on detection

Since prestimulus sensorimotor alpha amplitude modulated somatosensory perception, we hypothesized that alpha oscillations mediated the effect of cardiac phase on detection. To test this hypothesis, we calculated detection rates separately for systole and diastole trials within each alpha bin, where alpha amplitudes were comparable (*F*_1,36_ = 0.89, *p* = 0.35). Both cardiac phase and alpha levels significantly correlated with detection rate (within-subject ANOVA test, *F*_1,36_ = 15.82, *p* = 3·10^−4^ and *F*_2.93,105.30_ = 12.05, *p* = 1·10^−6^) but there was no significant interaction effect (*F*_4,144_ = 0.34, *p* = 0.85; **Fig. 5d**). This result indicated that detection rates differed between systole and diastole in the presence of comparable sensorimotor alpha amplitude levels.

To further test the relationship between prestimulus sensorimotor alpha amplitude, cardiac phase and detection, general linear mixed-effects modeling (GLMM) regressions were fitted at the single-trial level. Regressions that included only the cardiac phase or only the alpha amplitude were highly significant compared to a null model, i.e., a model with no relationship assumed (*cardiac model*: χ^2^ = 18.07, *p* = 4·10^−4^; *alpha_1_ model*: χ^2^ = 121.71, *p* = 2·10^−16^; **Supplementary Table 1**). The comparison of the *alpha_1_ model* and the *cardiac model* favored the *alpha_1_ model* (χ^2^ = 103.64, *p* = 2·10^−16^; **Supplementary Table 1**). The *additive_1_ model* that included both cardiac phase and alpha amplitude fitted the data significantly better than the *alpha_1_ model* (χ^2^ = 17.41, *p* = 0.002) and an *interaction_1_ model* that included an interaction between cardiac phase and alpha (χ^2^ = 1.51, *p* = 0.91; **Supplementary Table 1**). To illustrate the best model, the *additive_1_ model*, with numbers: If a stimulus was preceded by an alpha amplitude of 0.5 µV (1 standard deviation below the mean amplitude), the detection rates for stimuli in diastole and systole would be 56% and 53%, respectively. When prestimulus alpha amplitude increased to 1.4 µV (1 standard deviation above the mean amplitude), the detection rates for stimuli in diastole and systole would decrease to 49% and 46%, respectively. In summary, these results suggest that sensorimotor alpha and cardiac phase have independent effects on detection, i.e., alpha is not mediating the effect of cardiac phase on somatosensory detection.

### Prestimulus sensorimotor alpha does not mediate the effect of HEP on detection

To test whether prestimulus alpha amplitude mediated the relationship between HEP and detection, detection rates were calculated separately for low and high HEP levels within each alpha bin, where alpha amplitudes were similar between low and high HEP (*F*_1,36_ = 0.14, *p* = 0.71). A within-subject ANOVA showed significant main effects of both HEP (*F*_1,36_ = 38.71, *p* = 4·10^−7^) and alpha amplitude levels (*F*_4,144_ = 10.37, *p* = 2·10^−7^) for the detection rate with no significant interaction between them (*F*_4,144_ = 0.75, *p* = 0.56; **Fig. 5e**). This result shows that the HEP effect was additive to the effect of alpha levels on detection.

To confirm the additive effect of the amplitudes of prestimulus sensorimotor alpha and HEP on detection at the single-trial level, we calculated GLMM regression fits (cf. previous section). Regressions that included only alpha or only HEP, respectively, as predictors were highly significant compared with a null model (*alpha_2_ model*: (χ^2^ = 60.27, *p* = 5·10^−13^; HEP model: χ^2^ = 85.29, *p* = 2·10^−16^; **Supplementary Table 2**). The comparison of the *alpha_2_ model* and the *HEP model* favored the *HEP model* (χ^2^ = 25.02, *p* = 2·10^−16^; **Supplementary Table 2**). The *additive_2_ model* including both HEP and alpha amplitude in the regression fitted the data better than the *alpha_2_ model* (χ^2^ = 62.73, *p* = 1·10^−12^) and the *interaction_2_ model* (χ^2^ = 0.57, *p* = 0.45; **Supplementary Table 2**). To illustrate the best model, the *additive_2_ model*, with numbers: If a stimulus was preceded by a HEP amplitude of −1.7µV and an alpha amplitude of 0.5µV (1 standard deviation below the mean amplitude), the probability of detecting a stimulus was 59%. This probability would decrease to 51% if only the HEP amplitude would increase to 1.6μV and to 51% if only the alpha amplitude would increase to 1.4μV. If both HEP and alpha amplitudes would increase to 1.6μV and 1.4μV (one standard deviation above the mean amplitude), respectively, the detection probability would decrease to 43%. The GLMM results further support that sensorimotor alpha and HEP have independent effects on detection. Thus, alpha is also not mediating the effect of HEP on somatosensory detection.

### Controls for volume conduction effect

Moreover, we ascertained that the observed SEP differences between the two cardiac phases as well as the HEP effect on detection were not likely to be explained by differences in cardiac electrical activity, which might have caused differences in the EEG by volume conduction (Gray, Minati, Paoletti, & Critchley, 2010; Montoya et al., 1993; Park et al., 2014). First, we examined whether possible ECG artifacts were successfully eliminated during the calculation of SEP differences between systole and diastole (see Methods for further detail; **Supplementary Fig. 4a-c**): We tested whether the ECG waveform difference between the systole and diastole trials were canceled out after ECG artifact correction (**Supplementary Fig. 4d-f**). The comparison between two residual ECG waveforms for systole and diastole trials revealed no significant difference (no clusters were found, **Supplementary Fig. 4f**). Thus, the observed differences in SEP amplitudes between systole and diastole cannot be attributed to differences in cardiac electrical activity. Second, we checked whether the response to heartbeats preceding hits and misses differed in the ECG data. The ECG data looked similar for hits and misses (**Supplementary Fig. 5a**). The cluster statistics on the ECG data 296 – 400 ms after the R-peak did not show any significant difference between hits and misses (no clusters were found, **Supplementary Fig.5a**). Correcting the EEG data for the cardiac artifact using independent component analysis did not significantly change the results (**Supplementary Fig. 5b**). Therefore, HEP differences preceding hits and misses cannot be explained due to differences in cardiac electrical activity.

## DISCUSSION

We show that the timing of a somatosensory stimulus, with respect to the cardiac cycle, along with the amplitude of the prestimulus heartbeat-evoked potential (HEP) shape conscious perception and the somatosensory evoked potential (SEP). More specifically, detection rates were higher during diastole than systole and inversely related to the amplitude of the preceding HEP. Differential psychophysical effects of cardiac phase and HEP were observed on sensitivity and criterion, respectively. Furthermore, the cardiac phase influenced only late components of the SEPs (P300) whereas the effects of HEP amplitude were observed in both early (starting with P50) and late SEP components. While prestimulus alpha power also influenced perception and somatosensory processing, its effect was independent of both heartbeat-related effects on conscious perception, i.e., alpha power and heartbeat-related events had an additive impact on somatosensory perception.

Our first main finding, the modulation of perception and neural response along the cardiac cycle, seems best explained by periodical modulations of perception in a predictive coding framework, in which the brain is continuously producing and updating a model of sensory input. This model not only concerns exteroceptive stimuli but also interoceptive signals such as the heartbeat. Each heartbeat and its concomitant pulse wave lead to transient physiological changes in the entire body. These repeating cardiac fluctuations are treated as predictable events and attenuated by the brain to minimize the likelihood of mistaking these self-generated signals as external stimuli (Barrett & Simmons, 2015; Seth & Friston, 2016). Likewise, external stimuli coinciding with these predictable bodily events are inhibited. For instance, visual stimuli synchronous to the heartbeat have been shown to be suppressed compared with those presented asynchronously (Salomon et al., 2016). Of relevance for our study, heartbeat-related pressure fluctuations are tightly coupled with the firing pattern of afferent neurons in the fingers (Macefield, 2003). These neurons fire in response to the pressure wave that reaches its maximum after around 200 to 400 ms after the R-peak within systole (van Velzen, Loeve, Niehof, & Mik, 2017). We postulate that the same top-down mechanism, which suppresses the perception of heartbeat-related firing changes in afferent finger neurons (Macefield, 2003), also interferes with the perception of weak external stimuli to the fingers. This would only occur if presented during the same time period in systole – and more precisely between 200 and 400 ms after the R-peak. In line with this view, we further showed that individuals who had less heart rate variability, in other words, who should have better (temporal) predictions of the “next” heartbeat, showed higher perceptual suppression of stimuli during systole. Furthermore, this effect reflected changes in sensitivity, i.e., a weak input during systole is more likely to be regarded as pulse-associated “internal noise”, and thus the differentiation between the stimulation and “noise” becomes more difficult. This could also explain why localization becomes worse during systole.

A reduction of the P300 amplitude accompanied the cardiac-phase associated modulation in somatosensory perception and sensitivity during systole compared to diastole. As there were no differences in earlier SEP components, it seems rather unlikely that a peripheral mechanism explains the cardiac cycle effects on perception. If activity were reduced in the sensory receptors of the fingers during systole, it would yield a weaker bottom-up activity in the primary somatosensory cortex (SI) during systole and we would have observed SEP differences within 100 ms following the stimulus onset (Allison, McCarthy, & Wood, 1992). Since we found differences only for a later SEP component, the cardiac phase-related perceptual effects seem to be related to central neural processes (e.g., prediction). The P300 component has in fact been associated with predictive coding (Friston, 2005; Friston, Kilner, & Harrison, 2006) and has been suggested to be an indicator of conscious awareness (Auksztulewicz, Spitzer, & Blankenburg, 2012; Dehaene, Sergent, & Changeux, 2003; Sergent, Baillet, & Dehaene, 2005). Fittingly, the suppression of recurrent activity within the somatosensory network in the later stages of stimulus processing would be expected to reduce P300 amplitude (Auksztulewicz et al., 2012; Dehaene et al., 2003; Lamme, 2006). Taken together, the decreased P300 amplitude and lower sensitivity for somatosensory stimuli during systole might indicate a less efficient propagation of neuronal activity to higher processing levels (Vugt et al., 2018). In the context of the global neural workspace theory (Dehaene et al., 2003), decreased sensitivity prevents “ignition” of conscious perception of a stimulus by interfering with its processing within the higher-order sensory cortices. This prevents the broadcasting of the stimulus and therefore conscious perception of it.

Our second main finding links HEP amplitudes to the processing of weak somatosensory stimuli. Specifically, we show that HEP in the time range of 296 to 400 ms showed higher positivity for misses than hits over centroparietal electrodes. That is, the amplitude (positivity) of HEP was inversely related to detection as well as stimulus localization. This effect of HEP is not likely to reflect physiological differences in the functioning of the heart (a peripheral effect) since we could not detect any significant changes in cardiovascular measures (ECG, heart rate, heart rate variability) preceding hits or misses. Furthermore, in an SDT-based analysis, we have shown that the HEP effect was mainly related to changes in the criterion. With increasing HEP, participants adopted a more conservative bias for detection. A conservative bias has been shown to be associated with lower baseline firing rate across different brain regions, pushing neurons away from the threshold for “ignition” (Vugt et al., 2018). Supporting the mechanism of criterion, changing baseline firing rates in the brain, we found that the increasing prestimulus HEP amplitudes had a negative effect on the amplitude of both early (P50) and later SEP components (N140, P300). Similar modulation of early SEP components (P50) has previously been shown along with shifts of spatial attention (Forschack et al., 2017; Schubert et al., 2008). Given that HEP amplitude has been found to be significantly higher during interoceptive compared to exteroceptive attention (García-Cordero et al., 2017; Petzschner et al., 2019; Villena-González et al., 2017), we propose that the modulations of HEP amplitude reflect attentional shifts between external stimuli (exteroception) and internal bodily states (interoception). In line with this view, it has been suggested that the sudden ‘‘ignition’’ of a spontaneous internal activity can block external sensory processing (Dehaene & Changeux, 2005). Similarly, heartbeat-related signals, which have been suggested to contribute to spontaneously active and self-directed states of consciousness (Park et al., 2014), might prevent “ignition” of the upcoming somatosensory stimulus. Overall, the most plausible explanation for our findings seems to be that a shift from external to internal attention, reflected by HEP amplitude increases, interferes with conscious perception of external somatosensory stimuli by decreasing the baseline firing rates within the somatosensory network.

In the visual domain, a recent study also proposed that HEPs can predict the detection of weak stimuli (Park et al., 2014). Interestingly, Park et al. (2014) reported that larger heart-evoked activity measured using magnetoencephalography (MEG) was associated with better external perception, while we observed the opposite pattern. These differences might be due to the different sensory modalities tested, i.e., the allocation of attentional resources to interoception may vary for the detection of somatosensory and visual stimuli. In support of this notion, during a state of interoceptive attention, somatosensory cortex shows higher but visual cortex shows lower coupling to the anterior insular cortex, a key area for interoception (Wang et al., 2019). Furthermore, the somatosensory cortex has been indicated as one of the sources of HEPs (Kern et al., 2013; Pollatos, Kirsch, & Schandry, 2005) and as playing a substantial role for interoception (Critchley, Wiens, Rotshtein, Öhman, & Dolan, 2004; Khalsa, Rudrauf, Feinstein, & Tranel, 2009). Therefore, it seems plausible that heart-related processes in the interoceptive cortices, notably involving somatosensory but less so visual areas, may interfere differently with the processing of exteroceptive somatosensory and visual signals.

Our third main finding relates heartbeat-associated effects to ongoing neural activity. First, we attempted to confirm the influence of prestimulus sensorimotor alpha activity on somatosensory perception as shown in previous studies (Craddock, Poliakoff, El-deredy, Klepousniotou, & Lloyd, 2017; Jones et al., 2010; Schubert et al., 2009). We observed that during periods of weak prestimulus alpha amplitude, detection rates increased, which reflected a more liberal detection criterion. This finding is consistent with studies in the visual (Iemi et al., 2017) and somatosensory domain (Craddock et al., 2017). Even though detection has already been associated with lower alpha levels (Jones et al., 2010; Schubert et al., 2009; van Dijk et al., 2008), the relationship between somatosensory localization and alpha amplitudes – to the best of our knowledge – has not been reported so far. In the visual domain, when localization and detection tasks were tested with a block design, detection but not localization was shown to vary across alpha levels (Iemi et al., 2017). For the somatosensory domain, we showed that not only detection rates but also localization rates increased with decreasing prestimulus alpha amplitudes. Given the effect of alpha on somatosensory perception, we tested whether sensorimotor alpha oscillations modulated the heartbeat-related effects on detection. Our analysis showed that neither of the two heartbeat-related effects on perception (i.e., the cardiac phase and the HEP amplitude) was mediated by pre-stimulus alpha amplitude, but rather both are independent and additive to the effect of prestimulus sensorimotor alpha amplitude.

Several pathways relating cardiac activity to the brain have been suggested. Most notably, baroreceptor activation might inform cortical regions about timing and strength of each heartbeat (Critchley & Garfinkel, 2018). Baroreceptors are maximally activated during systole and their stimulation has been suggested to reduce cortical excitability (Rau & Elbert, 2001). Thus, the systolic activation of baroreceptors might inform predictive mechanisms in the brain concerning when to attenuate the processing of heartbeat-coupled signals. Other than through baroreceptors, cardiac signals might also reach the cortex through direct projections of cardiac afferent neurons to the brain (Park & Blanke, 2019) or via somatosensory afferents on the skin (Khalsa et al., 2009). While presently it is not clear which of these pathways is most relevant for heart-brain interactions, our results are consistent with the notion of the somatosensory cortex as an important relay center for cardiac input (Critchley et al., 2004; Kern et al., 2013; Khalsa et al., 2009; Pollatos et al., 2005). How this relay center modulates the relationship between interoception and exteroception is an interesting topic for future research.

In conclusion, timing of stimulation along the cardiac cycle and spontaneous fluctuations of HEP amplitudes modulate access of weak somatosensory stimuli to consciousness and induce differential effects on somatosensory-evoked potentials. We explain these fundamental heart-brain interactions within the framework of interoceptive predictive coding (stimulus timing) and spontaneous shifts between interoception and exteroception (HEP amplitudes). These findings on heartbeat-related perceptual effects might serve as an example how in general body-brain interactions can shape our cognition.

## MATERIALS AND METHODS

### Participants

Forty healthy volunteers were recruited from the database of the Max Planck Institute for Human Cognitive and Brain Sciences, Leipzig, Germany. Three subjects were excluded from the analysis due to technical problems during the experiment. Data from thirty-seven subjects were analyzed (twenty females, age: 25.7 ± 3.9 years (mean ± s.d.), range: 19-36 years). Some experimental blocks were excluded from the data analysis due to data acquisition failures (8 blocks from 5 subjects), false alarm rates > 40 % (8 blocks from 8 subjects), responding with the wrong finger in the task (4 blocks from 3 subjects) and observation of closed eyes during the task (3 blocks from 1 subject). After these exclusions, a total of 274 experimental blocks with 32880 trials in 37 subjects were analyzed. The study was approved by the Ethical Committee of the University of Leipzig’s Medical Faculty (No: 462-15-01062015). All subjects signed written informed consent and were paid for their participation.

### Somatosensory stimulation and task design

Electrical finger nerve stimulation was performed with a constant-current stimulator (DS5; Digitimer) using single square wave pulses with a duration of 200 μs. Steel wire ring electrodes were placed on the middle (anode) and the proximal (cathode) phalanx of the index and the middle finger of the left hand, respectively.

In the experiment, participants performed a yes/no detection and a two-alternative forced choice localization task on every trial. At the beginning of each trial, a black dot appeared on the screen for 600 ms. Participants then expected to get stimulation on either the index or the middle finger of their left hand. 600 ms after the stimulation, participants “were asked” (via “*yes/no?”* on the screen) to report as quickly as possible *whether* they felt a stimulus on one of their fingers or not. They responded “yes” if they felt the stimulus and “no” if not by using their right index finger. Thereafter, participants were asked to answer *where* the stimulation has occurred. They were explicitly told “to guess” even if they reported not feeling the stimulus in the first question. If they located the stimulus on the left index finger, they were asked to use their right index finger to answer and to use their right middle finger if they located the stimulus on the left middle finger. The next trial started immediately after responding to the localization question. In total, every participant completed eight blocks. Each block contained 100 trials with electrical stimulation (50 trials for each finger) and 20 trials without any stimulation (catch trials). The duration of each block was ∼8 minutes. To find stimulus intensities with 50 % detection probability (i.e., threshold), we applied a two-step procedure before starting the experiment: First, we roughly estimated the lowest stimulus intensity for which participants could report a sensation by applying the method of limits with ascending intensities separately for the index and the middle finger (Forschack et al., 2017; Taskin, Holtze, Krause, & Villringer, 2008). Second, we used a yes/no detection task (as described above) containing catch trials and 6 stimulus intensities around this predicted stimulus intensity (15% below, identical to, 20%, 40%, 60% and 80 % above) for each finger. The 50% threshold intensity for each finger was estimated from the participant’s psychometric function (Ehrenstein & Ehrenstein, 1999). To control for threshold stability, stimulus intensities were readjusted after each block.

*Hit*, *miss*, *false alarm* (FA) and *correct rejection* (CR) terms were calculated for the yes/no detection task in this study. A *hit* was reporting the presence of a stimulus when it was present; a *miss* was reporting the absence of a stimulus even though it was present. For catch trials (i.e., no stimulus was presented), an *FA* was reporting the presence of a stimulus, while a *CR* was reporting its absence. The terms *correct localization* and *wrong localization* were used to describe the localization task performance. *Correct localization* was reporting the stimulus location correctly; *wrong localization* was reporting it incorrectly.

### Recordings

EEG was recorded from 62 scalp positions distributed over both hemispheres according to the international 10–10 system, using a commercial EEG acquisition system (actiCap, BrainAmp, Brain Products). The mid-frontal electrode (FCz) was used as the reference and an electrode placed on the sternum as the ground. Electrode impedance was kept ≤ 5 k∧ for all channels. EEG was recorded with a bandpass filter between 0.015 Hz and 1 kHz and digitized with a sampling rate of 2.5 kHz. An ECG electrode connected to the EEG system was placed under the participant’s left breast to record the heart activity.

### Data analysis

We applied two complementary approaches – circular and binary analysis – to examine detection and localization across the cardiac cycle (Kunzendorf et al., 2019). For these analyses, we first extracted the R-peaks from the ECG data by using Kubios HRV Analysis Software 2.2 (The Biomedical Signal and Medical Imaging Analysis Group, Department of Applied Physics, University of Kuopio, Finland) and visually corrected for inaccurately determined R-peaks (<0.1%). From RR interval time series during the whole experiment, we calculated the standard deviation of RR intervals (SDNN) and natural-log transformed SDNN values to calculate heart rate variability (Shaffer & Ginsberg, 2017; Tarvainen, Niskanen, Lipponen, Ranta-aho, & Karjalainen, 2014).

### Circular analysis

We tested detection and localization over the entire cardiac cycle, from one R-peak to the next one, by using circular statistics (Pewsey, Neuhäuser, & Ruxton, 2013). We calculated the relative position of the stimulus onset within the cardiac cycle with the following formula

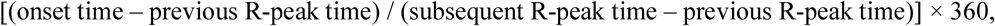

which resulted in values between 0 and 360 degrees (0 indicating the R-peak before stimulus onset). The distribution of stimulus onsets was tested individually for each participant with a Rayleigh test for uniformity. Two participants were excluded from further circular analyses due to non-uniformly distributed stimulation onsets across the cardiac cycle, (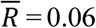, *p* = 0.04; 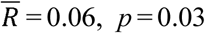). For the rest of the participants (N=35), the assumption of uniform onset distributions was fulfilled. We calculated the mean phase value at which different performances occurred (detection task: hit and miss; localization task: correct localization and wrong localization) for each participant. At the group level, it was tested whether the distribution of a specific performance score (e.g., hits) deviated from the uniform distribution with Rayleigh tests (Pewsey et al., 2013). The Rayleigh test depends on the mean vector length out of a sample of circular data points and calculates the mean concentration of these phase values around the circle. A statistically significant Rayleigh test result indicates the non-uniform distribution of data around the circle, that is the cardiac cycle.

### Binary analysis

Detection and localization performances were examined across the systolic and diastolic phases of the cardiac cycle. We defined *systole* as the time between the R-peak and the end of the t-wave (Motyka et al., 2019). We used the systolic length of each cardiac cycle to define *diastole* as a diastolic window of equal length placed at the end of the cardiac cycle. The equal length of systole and diastole was used to equate the probability of having a stimulus onset in the two phases of the cardiac cycle. To determine the end of t-wave, a trapez area algorithm was applied in each trial (Vázquez-Seisdedos, Neto, Marañón Reyes, Klautau, & Limão de Oliveira, 2011). This method has advantages compared to an approach with fixed bins (e.g., defining systole as the 300-ms time window following the R-peak) because it accounts for within- and between-subject variations in the length of systole and diastole (i.e., the heart rate). The results of the automated algorithm were visually quality-controlled. 27 trials for which the algorithm failed to calculate t-wave end and produced an abnormal systole length (more than 4 standard deviations above or below the participant-specific mean systole) were removed from further binary analyses. Mean systole (and diastole) length obtained from these analyses was 333 ± 21ms. Each trial was categorized depending on whether the stimulus occurred during systole or diastole. The average number of trials categorized as systole was 338 ± 51 and as diastole was 342 ± 59.

### Data preprocessing

EEG and ECG data were analyzed offline using EEGLAB (Delorme & Makeig, 2004) and FieldTrip (Oostenveld, Fries, Maris, & Schoffelen, 2011) toolbox algorithms as well as custom-built scripts on a Matlab platform (Mathworks Inc.). An anti-aliasing filter with a 112.5 Hz-cutoff was used before downsampling individual datasets to 250 Hz. After all blocks were concatenated, data were first high-pass filtered with 0.5 Hz and then low-pass filtered with 45 Hz using a 4^th^ order of Butterworth filter. The EEG channels that had a flat line longer than five seconds or showed less than 85% correlation with its reconstructed activity from other channels were removed and interpolated using their neighboring channels. After a principal component analysis was applied, data underwent an independent component analysis (ICA) using an extended infomax algorithm to remove sources of heartbeat, ocular and muscle artifacts (Delorme, Palmer, Onton, Oostenveld, & Makeig, 2012; Li, Ma, Lu, & Li, 2006). ICA components with cardiac field artifact were determined by segmenting ICA components depending on the R-peak of the ECG electrode and visually selecting the components whose activities were matching the time course of R-peak and t-wave of the ECG. After removing artifactual ICA components, the artifact-free components were forward-projected for the subsequent analysis steps. Afterward, the data were re-referenced to the average reference.

### Somatosensory-evoked potentials

Data were segmented from −1000 to 2000 ms with respect to stimulus onset separately for trials where the stimulation occurred during systole vs diastole. After segmenting data, we performed baseline correction using 100 to 0 ms prestimulus window. Testing for the maximum positive deflection of the early SEP component P50 (40 to 60 ms) showed that the right primary somatosensory area, contralateral to the stimulated hand (Zhang & Ding, 2010), was represented by the C4 electrode. Therefore, the statistical analysis of SEP amplitude was performed on the C4 electrode (Nierhaus et al., 2015). To cancel out possible effects of blood circulation, we estimated the cardiac artifact in the EEG data. For this purpose, random triggers were placed over cardiac cycles outside the stimulation window-see **Fig. 1**). Then, we classified the arbitrary triggers as systole or diastole depending on the position of the trigger in the cardiac cycle. After the classification, data were segmented around these triggers (−1000 to 2000 ms) and averaged separately for systole and diastole to estimate the cardiac artifact during systole and diastole for each EEG channel per subject. We baseline-corrected these signals 100 ms before the onset of the arbitrary triggers (**Supplementary Fig. 4**). To prevent any possible ECG-induced artifact on the SEPs, we subtracted the mean systolic and diastolic artifacts from the SEPs during systole and diastole trials, respectively (Gray et al., 2010).

### Heartbeat-evoked potentials

After preprocessing data as described above, we selected the cardiac cycles containing a stimulus. We only chose the trials in which the stimulus onset was at least 400 ms after the preceding R-peak (corresponding to diastole). We determined HEPs by segmenting the preprocessed EEG data from −1000 to 2000 ms around the R-peak separately for hits and misses as well as for correct localizations and wrong localizations. In this way, we could calculate the prestimulus HEPs, which have been reported between 250 - 400 ms after the R-peak (Kern et al., 2013; Schandry et al., 1986; Schandry & Weitkunat, 1990).

### Time-frequency analyses

We performed time-frequency analyses to investigate sensorimotor alpha activity locked to stimulus onset. For sensorimotor alpha, we selected ICA components representing sensorimotor rhythms to eliminate effects of the occipital alpha activity as described previously by our group (Forschack et al., 2017; Nierhaus et al., 2015). 1 – 7 components per participant (mean 3± 1 SD) were selected and included in the analysis of somatosensory oscillatory activity. Then, data were segmented (−1000 to 2000 ms), ECG-induced artifacts for systole and diastole were calculated, and subtracted from the data as described in the previous section. Wavelet analysis was performed for frequencies from 5 to 40 Hz in 1 Hz increments to allow for a time-resolved frequency analysis of event-related power modulation. The wavelet transformation was performed on every trial using wavelet cycle lengths from 4 to 10 cycles increasing with frequency in linear steps. Then, the time-frequency response was averaged for systole and diastole conditions. We focused on the effects of prestimulus alpha activity in our statistical analysis to test whether the perceptual effect of the cardiac cycle on detection is influenced by prestimulus oscillatory activity (−300 to 0 ms) over contralateral somatosensory area.

### Analyses according to Signal Detection Theory (SDT)

Sensitivity (d’) and criterion (c, response bias) were calculated according to SDT (Macmillan & Creelman, 2004): d’ and c were calculated as z(HR)-z(FAR) and −[z(HR)+z(FAR)]/2, respectively, with HR corresponding to hit rate and FAR corresponding to false alarm rate. A log-linear correction was used to compensate for extreme false alarm proportions(Hautus, 1995) since two of the thirty-seven participants produced no false alarms. Localization d’ prime was calculated as √2 * z (correct localization rate).

### Statistical analyses

Assessment of statistical significance for “two-condition comparisons” in EEG data was based on cluster-based permutation *t*-tests as implemented in the FieldTrip toolbox (Maris & Oostenveld, 2007; Oostenveld et al., 2011). In this procedure, adjacent spatio-temporal or spatio-spectro-temporal points for which *t*-values exceed a threshold are clustered (cluster threshold *p*-value: 0.05). Then the cluster statistics are calculated by taking the sum of *t*-values of all points within each cluster. The type I error rate was controlled by evaluating the cluster-level statistics under a randomized null distribution of the maximum cluster-level statistics. To determine the distribution of maximal cluster-level statistics obtained by chance, condition labels were randomly shuffled 1000 times. For each of these randomizations, cluster-level statistics were computed and the largest cluster-level statistic was entered into the null distribution. Finally, the experimentally observed cluster-level statistic was compared against the null distribution. Clusters with a *p*-value below 0.05 (two-tailed) were considered “significant”. We expected to observe differences in SEPs over contralateral somatosensory cortex indexed by C4 electrode. Therefore, in the comparisons of somatosensory related activity, we only used cluster statistics to test whether two experimental conditions differed in time over contralateral somatosensory cortex. In contrast, we did not *a priori* define a spatial region for HEP analyses but expected to observe a HEP between 250 and 400 ms after the R-peak (Kern et al., 2013; Schandry et al., 1986; Schandry & Weitkunat, 1990).

General linear mixed-effects modeling (GLMM) was used for mediation analyses since they both acknowledge both between- and within-participant variations in the data from the model’s fixed-effect estimates. GLMM was conducted in R (R Core Team, 2014) within the lme4 framework (Bates, Mächler, Bolker, & Walker, 2015). The models were defined in the following form: outcome ∼ predictor(s) + (predictor(s) | subject), which fits predictors of the fixed effect part (next to the “∼”) and predictors of the random effects part (in brackets) grouped by a factor, for which the predictors vary randomly, in our case, subjects.

First, we used GLMM to test whether the cardiac phase effect on detection was mediated by the prestimulus alpha amplitude. We computed five GLMMs regressing detection outcome (hit or miss): (1) one null model assuming no relationship, i.e., only the intercept served as predictor; (2,3) two models including either cardiac phase or alpha amplitudes as the fixed and random effect to regress detection; (4) an additive model regressing detection on both cardiac phase and alpha amplitude and (5) an interactive model assuming an interaction between cardiac phase and alpha amplitude to regress detection (see **Supplementary Table 1**). Second, we used GLMM to test whether prestimulus alpha mediated the effect of HEP on detection. We computed five GLMMs regressing detection outcome: (6) one null model; (7,8) two models including either HEP or alpha amplitudes as the fixed and random effect to regress detection; (9) an additive model regressing detection on both HEP and (10) alpha amplitude and an interactive model assuming an interaction between these two predictors (see **Supplementary Table 2**). In all the models containing alpha as a predictor, we used natural logarithmic transformation of alpha amplitude to normalize its distribution. To determine the best GLMM model explaining the data, maximum-likelihood ratio test statistics, which account for model complexity, were used.

If the sphericity assumption was violated in within-subject ANOVA, Greenhouse-Geisser correction was applied. All statistical tests were two-sided.

## ACKNOWLEDGEMENTS

We thank Luke Tudge, Mike X. Cohen, Sylvia Stasch, Dan J. Cook, Luca Iemi, Juan R. Loaiza, Stella Kunzendorf and Megan Peters for their valuable comments on the manuscript.

## AUTHOR CONTRIBUTIONS

E.A., F.I and A.V. designed the experiment, E.A. and F.I acquired the data. E.A., A.V., V.N., N.F., T.N. analyzed the data. E.A., N.F., T..N., M.G., M.G., P.M., V.N., A.V. wrote the paper.

