## Supplementary Files for "Heart-Brain Interactions Shape Somatosensory Perception and Evoked Potentials"

### **SUPPLEMENTARY DATA**

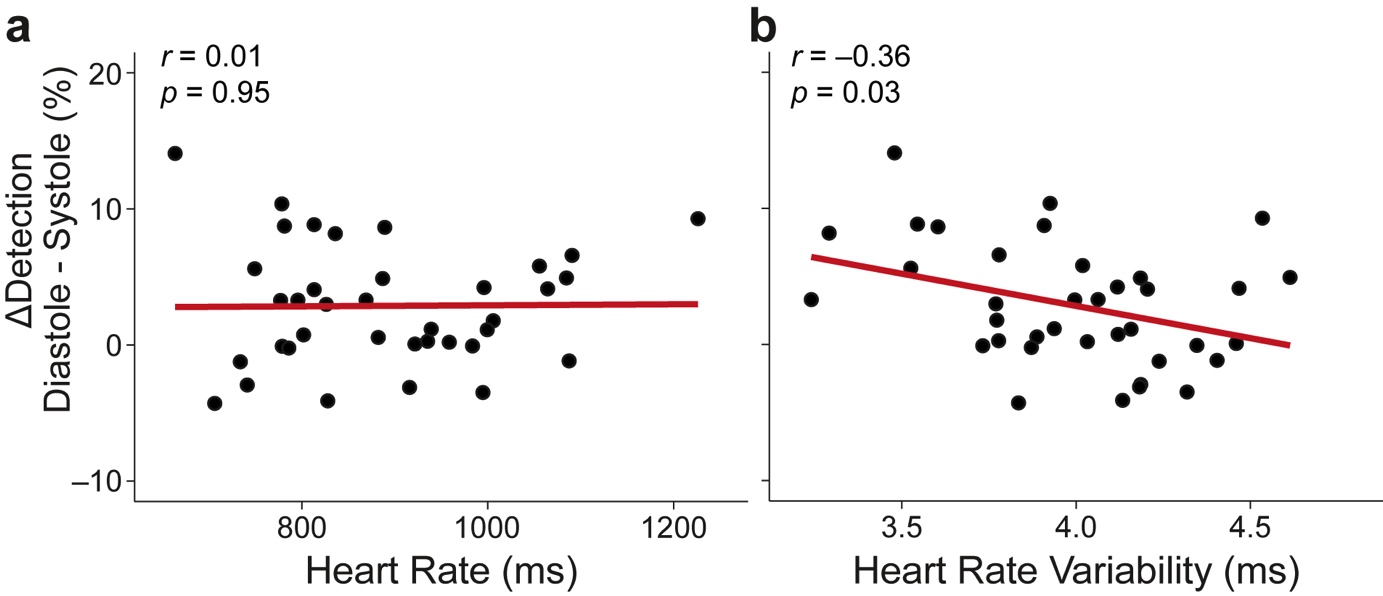

**Supplementary Figure 1** The change of detection between diastole and systole and its correlation with heart rate and heart rate variability. (**a**) Heart rate of subjects did not significantly correlate with their detection performance change between diastole and systole (Pearson’s correlation, *r*= 0.01, *p*=0.95) (**b**) Heart rate variability (i.e., the standard deviation of RR intervals, SDNN) of subjects negatively correlated with the change of detection performance between systole and diastole (*r*= -0.36, *p*=0.03).

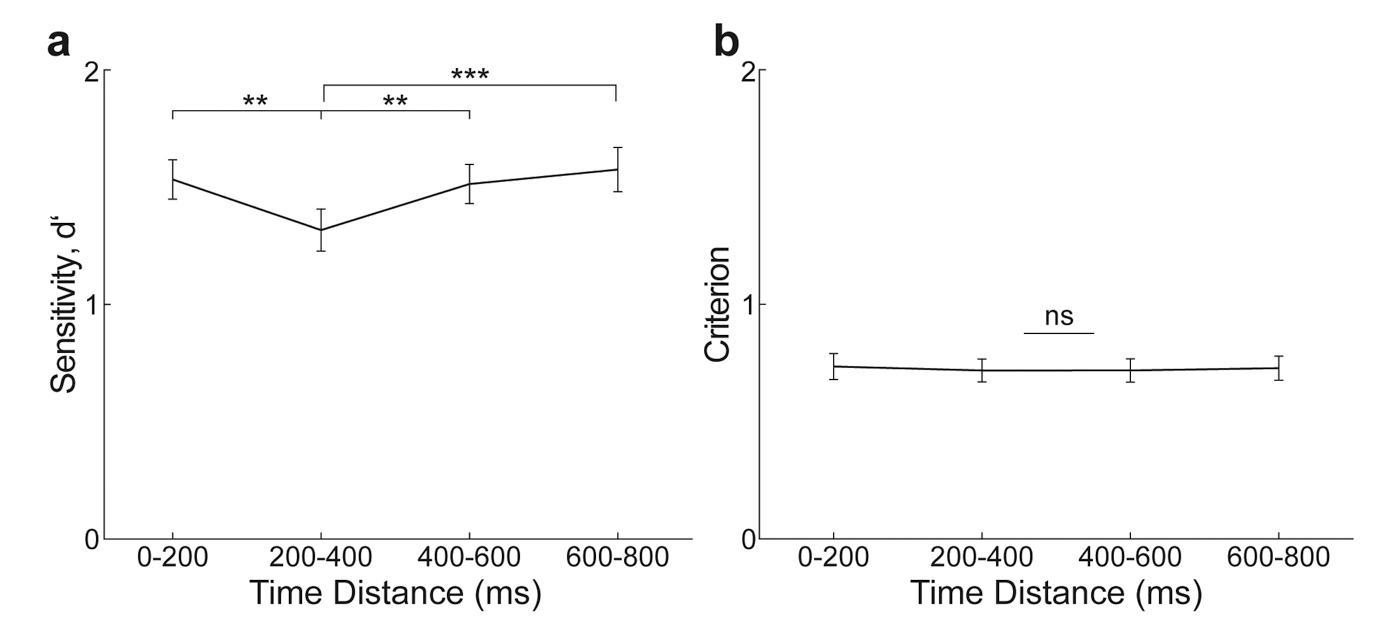

**Supplementary Figure 2** Sensitivity and criterion across four time windows of stimulus onset relative to the previous heartbeat (R-peak). (**a**) The detection sensitivity (d’) was lowest 200–400ms after the R-peak (*post-hoc* paired *t-*test between 0–200 and 200–400ms, *t*_36_=2.83, *p*=0.008 and between 200–400 and 400–600ms, *t*_36_=-3.48, *p*=0.001) (**b**) Criterion did not differ significantly between the four time windows (main effect of time, *F*_3,108_=0.10, *p*=0.96). Error bars represent SEMs. ***p*<0.005, ****p*<0.0005. ns, not significant.

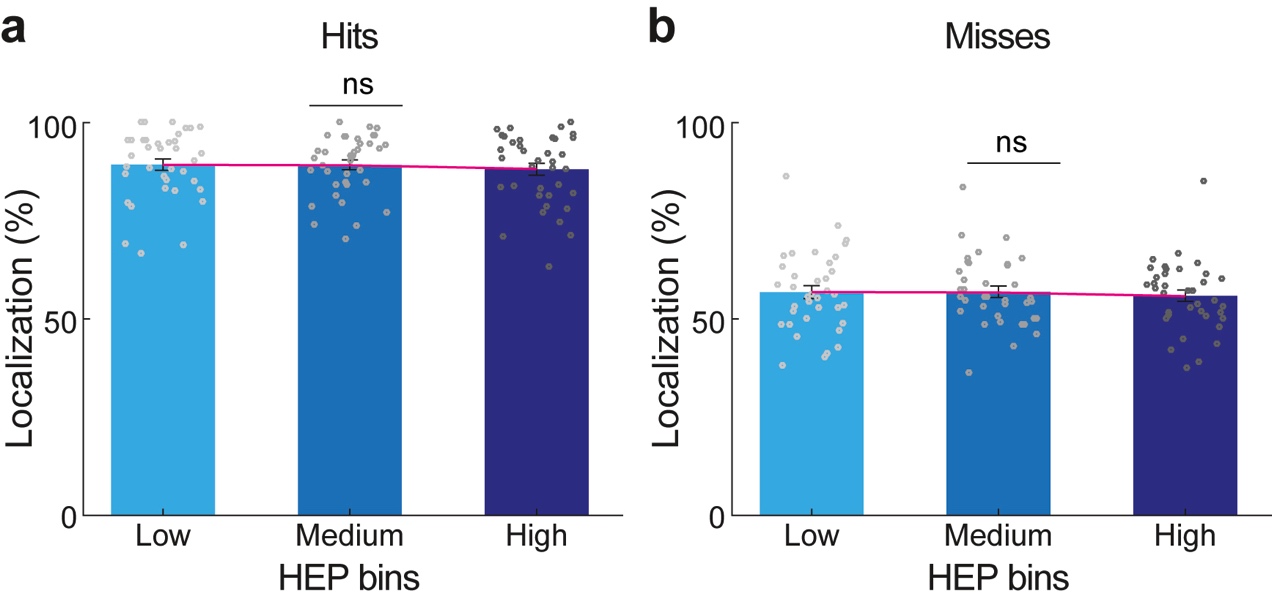

**Supplementary Figure 3** Correct localization of hits and misses across heartbeat-evoked potential (HEP) bins. (**a**) Correct localization of hits did not significantly change across increasing levels of HEP (within-subject ANOVA, *F*_2,72_=1.26, *p*=0.29) (**b**). Correct localization of misses did not significantly vary across HEP bins (*F*_2,72_=0.28, *p*=0.76). Error bars represent SEMs. ns, not significant.

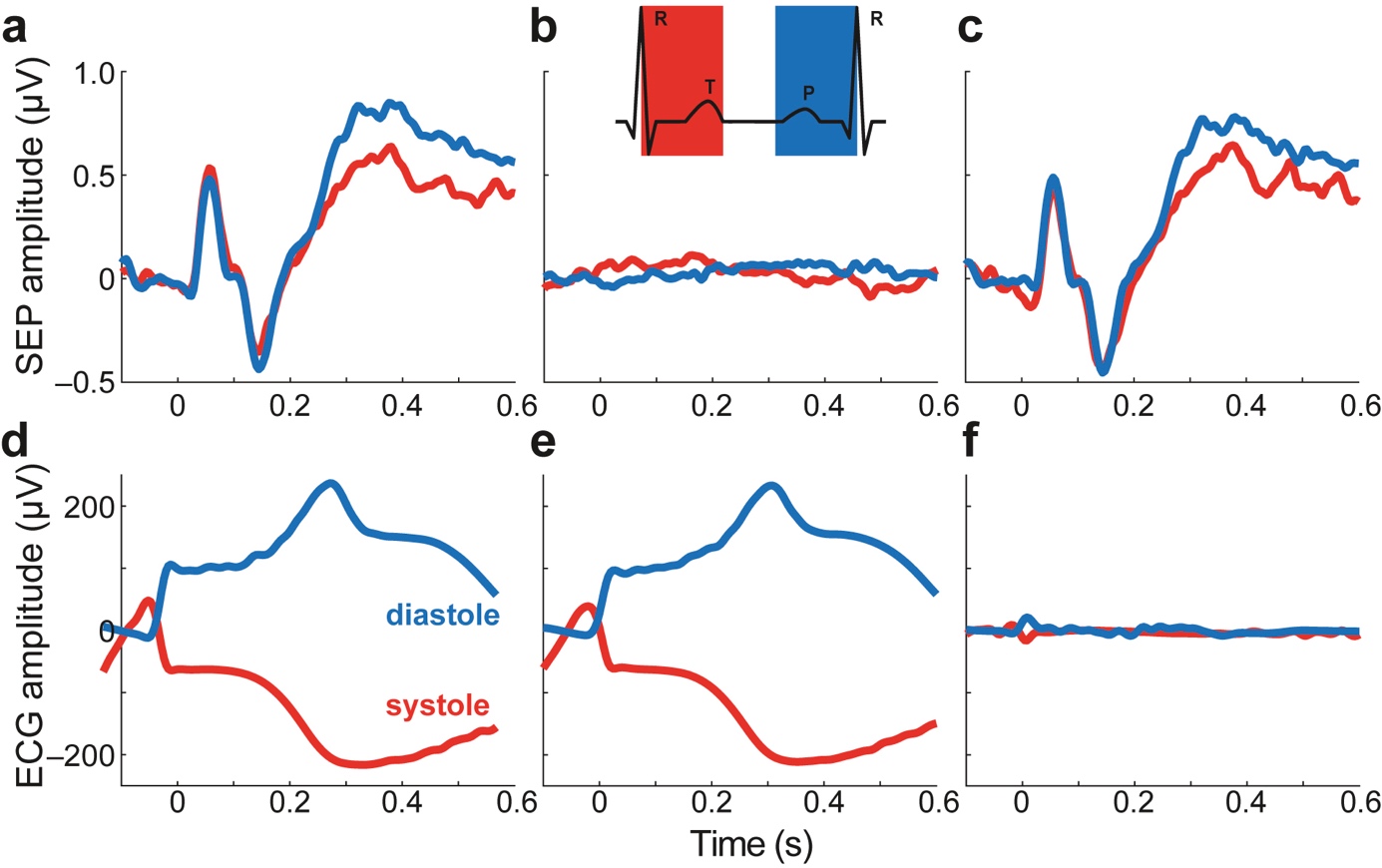

**Supplementary Figure 4** Effect of ECG artifact correction on stimulus-onset locked somatosensory-evoked potential (SEP) and electrocardiogram (ECG) amplitude (0=stimulation onset). (**a**) SEP at C4 before the artifact removal. (**b**) To cancel out the possible effects of ECG artifact, we estimated the cardiac artifact in the evoked responses by first placing random triggers along those cardiac cycles outside the stimulation window of the experiment). Then, we classified the arbitrary triggers as systole or diastole depending on the position of the trigger in the cardiac cycle. After the classification, we segmented data around the triggers and calculated the average cardiac artifact separately for systole and diastole in C4 electrode. (**c**) SEP during systole and diastole after the estimated artifact removal. The average estimation of the cardiac artifact for systole and diastole were subtracted from the SEP separately during systole and diastole. (**d**) Stimulus onset-locked ECG, grand average across participants before the artifact removal. (**e**) The estimated average cardiac artifacts on ECG amplitude relative to random triggers placed along the cardiac cycles excluding the stimulation window. (**f**) After the subtraction of the estimated cardiac artifact from the stimulus onset-locked ECG activity separately for systole and diastole, the difference in ECG amplitude during diastole versus systole is negligible. This analysis shows that the observed SEP differences between diastole and systole after ECG correction cannot be attributed to differences in cardiac electrical activity.

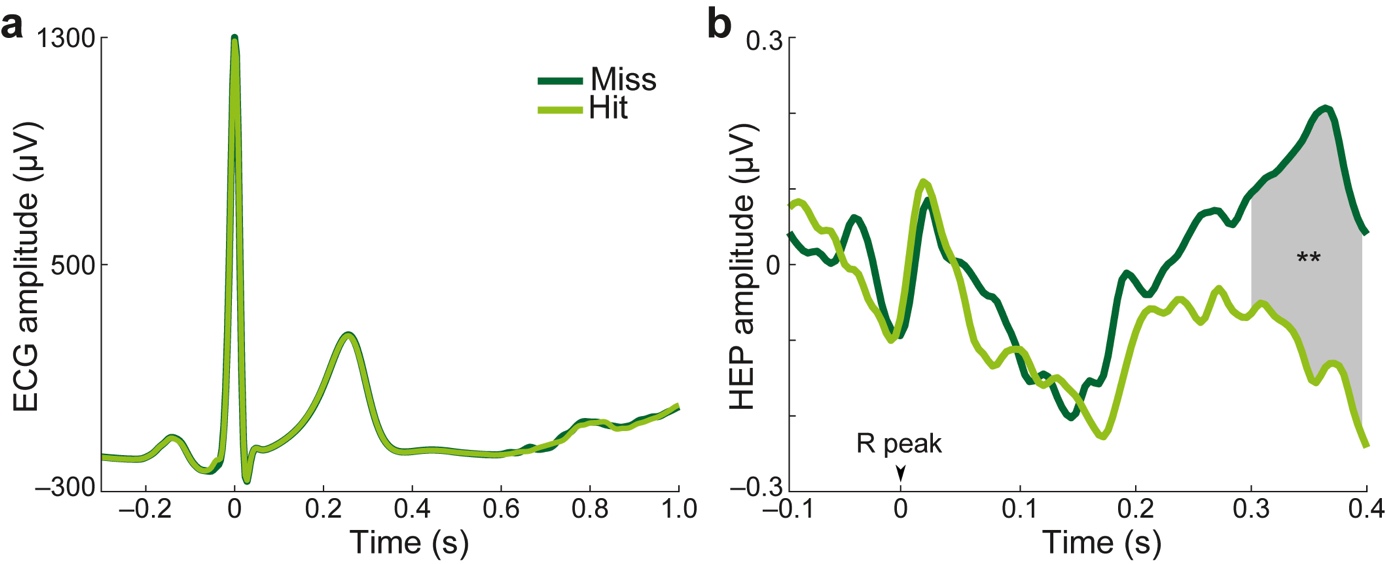

**Supplementary Figure 5** The difference of heartbeat-evoked potential (HEP) amplitude between hits and misses is not due to a cardiac field artifact (**a**) R-peak locked electrocardiogram (ECG) grand average across participants. We did not find any significant difference in ECG data between hits and misses (**b**) The HEP across cluster electrodes before cardiac field artifact removal with independent component analysis. The significant difference of HEP between hits and misses between 296-400ms (gray area) after the R-peak was conserved. ***p*<0.005.

**Supplementary Table 1 General linear mixed-effects modeling (GLMM) testing the relationship between prestimulus alpha amplitude, cardiac phase, and detection (model no. 1-5)**

| Model name | Glmer syntax | Likelihood | LRT |
| --- | --- | --- | --- |
| 1 – null_1_ | detection ~ 1 + (1 \| subject) | -14,165 |  |
| 2 – cardiac | detection ~ cardiac + (cardiac \| subject) | -14,156 | (1) $\chi2=18.07$*** |
| 3 – alpha_1_ | detection ~ alpha + (alpha \| subject) | -14,104 | (1) $\chi2=121.71$***  (2)$\chi2=103.64$*** |
| 4 – additive_1_ | detection ~ cardiac + alpha + (cardiac + alpha \| subject) | -14,096 | (3)$\chi2=17.41$** |
| 5 – interaction_1_ | detection ~ cardiac * alpha + (cardiac * alpha \| subject) | -14,095 | (4)$\chi2=1.51$ |

Likelihood shows the log-transformed likelihood of the models. Higher values of likelihood make the model more likely. LRT is the maximum likelihood ratio test comparing two models for the same dataset. More complex models (with more parameters) are compared with respective smaller ones, which gives a *χ2* and *p*-value. **p*< 0.05, ***p*< 0.005, *** *p*< 0.0005

**Supplementary Table 2 General linear mixed-effects modeling (GLMM) testing the relationship between prestimulus sensorimotor alpha amplitude, heartbeat-evoked potential, and detection (model no. 6-10)**

| Model name | Glmer syntax | Likelihood | LRT |
| --- | --- | --- | --- |
| 6 – null_2_ | detection ~ 1 + (1 \| subject) | - 10,007 |  |
| 7 – alpha_2_ | detection ~ alpha + (alpha \| subject) | - 9,976 | (6) $\chi2=60.27$*** |
| 8 – HEP | detection ~ HEP + (HEP \| subject) | - 9,964 | (6) $\chi2=85.29$***  (7)$\chi2=25.02$*** |
| 9 – additive_2_ | detection ~ HEP + alpha + (HEP + alpha \| subject) | - 9,933 | (8)$\chi2=62.73$*** |
| 10 – interaction_2_ | detection ~ HEP * alpha + (HEP * alpha \| subject) | - 9,932 | (9)$\chi2=$ 0.57 |

Models are evaluated as in Table 1. **p*< 0.05, ***p*< 0.005, *** *p*< 0.0005.
